# Automatic multisensory integration follows subjective confidence rather than objective performance

**DOI:** 10.1101/2023.06.07.544029

**Authors:** Yi Gao, Kai Xue, Brian Odegaard, Dobromir Rahnev

## Abstract

It is well known that sensory information from one modality can automatically affect judgments from a different sensory modality. However, it remains unclear what determines the strength of the influence of an irrelevant sensory cue from one modality on a perceptual judgment for a different modality. Here we test whether the strength of multisensory impact by an irrelevant sensory cue depends on participants’ objective accuracy or subjective confidence for that cue. We created visual motion stimuli with low vs. high overall motion energy, where high-energy stimuli yielded higher confidence but lower accuracy in a visual-only task. We then tested the impact of the low- and high-energy visual stimuli on auditory motion perception. We found that the high-energy visual stimuli influenced the auditory motion judgments more strongly than the low-energy visual stimuli, consistent with their higher confidence but contrary to their lower accuracy. A computational model assuming common principles underlying confidence reports and multisensory integration captured these effects. Our findings show that automatic multisensory integration follows subjective confidence rather than objective performance and suggest the existence of common computations across vastly different stages of perceptual decision making.

## Introduction

In daily life, individuals frequently receive information from multiple sensory modalities simultaneously. A substantial literature has examined how information from different sensory cues is automatically combined within sensory areas to form a single perceptual judgment^1–6^. Automatic multisensory integration occurs even when participants are explicitly informed that certain sensory cues are irrelevant to the task at hand^4,7–11^. Findings from this literature show that when sensory cues from one modality are particularly accurate or exhibit less noise, they exert a greater influence on perceptual judgments^2,3,5^.

A separate literature has examined people’s ability to metacognitively estimate how noise in the sensory input (e.g., due to low stimulus contrast, low motion coherence, or image blur) impacts accuracy via confidence ratings^12–15^. This literature has demonstrated that confidence is typically higher for stimuli that exhibit less stimulus noise^16,17^. However, many studies have also found that ^18–20^confidence and accuracy can often be dissociated such that participants give higher confidence in one condition compared to another even if accuracy for the two conditions is matched^21–25^. This raises the question: Does multisensory integration follow objective performance or subjective confidence when the two conflict?

Two competing hypotheses can be formulated. According to Hypothesis 1, multisensory integration follows objective accuracy, with metacognitive confidence having no impact. This view is motivated by extensive literature showing that multisensory integration is often an automatic process^11,26–29^, though it can be influenced by various factors, including attention, spatial and temporal alignment and physical parameters of the stimuli^30–33^. In contrast, metacognitive judgments of confidence are supported by networks involving the prefrontal cortex and other brain regions, allowing for both automatic and controlled processes that may rely on heuristics and other high-level cognitive mechanisms^34–37^. Hypothesis 1 would therefore predict that the visual stimulus with lower performance will have a smaller influence on multisensory integration regardless of its higher confidence. Conversely, according to Hypothesis 2, multisensory integration follows metacognitive confidence. In this view, participants need to first engage in a process of estimating the objective reliability of each stimulus in multisensory tasks. This estimation process is fallible. Hypothesis 2 would therefore predict that the visual stimulus with higher confidence will have a larger influence on the multisensory decision regardless of its lower accuracy.

Here, we adjudicated between these two competing hypotheses by investigating whether the strength of multisensory impact by an irrelevant sensory cue depends on participants’ objective accuracy or subjective confidence for the irrelevant cue. We employed an established method^21^ to induce a confidence-accuracy dissociation by manipulating the energy levels in a random-dot kinematogram. We found that the high-energy visual stimuli led to lower accuracy but higher confidence than low-energy visual stimuli in a visual-only task. Critically, we presented visual and auditory in congruent and incongruent directions across different trials and examined the automatic influence of high- and low-energy visual motion stimuli on auditory motion perception (leftward vs. rightward motion). We found strong support for Hypothesis 2. Specifically, consistent with their associated higher confidence but contrary to their lower accuracy, the high-energy stimuli had a larger impact on auditory perception. A simple computational model that assumes unified computations underlying multisensory integration and confidence reports reproduced this pattern of results. Our findings indicate that automatic multisensory integration follows subjective confidence rather than objective performance, suggesting the presence of shared computational processes across various stages of perceptual decision-making.

## Materials and Methods

### Participants

We collected data from 99 participants (30 females and 69 males, aged from 17 to 33, mean age = 20.5, SD = 2.6). Gender information was collected based on self-report. Participants’ race was collected but was not analyzed as it was irrelevant to the purpose of the study. The sample size for each group is sufficient to detect an effect, as calculated using G*Power (version 3.1.9.7), with the power set at 0.8, alpha at 0.05, and an effect size of 0.6 for a two-tailed paired t-test. A total sample size of 99 participants is expected to yield robust experimental results. We utilized a convenience sampling strategy. Participants were recruited from the student body at Georgia Tech and were compensated with either 1 SONA credit or monetary reward ($10/hr). While the sample may not fully represent the broader population, no significant differences are assumed between college students and other potential subject groups. All participants had normal or corrected-to-normal vision and normal hearing abilities. Before the experiment, all participants provided written consent approved by the Institutional Review Board of Georgia Institute of Technology (Protocol number: H21041).

### Stimuli

The experiment featured both visual and auditory motion stimuli. The visual stimuli were random-dot kinematograms that included three types of dot motion: (1) dots moving in the dominant direction, (2) dots moving in the non-dominant direction (opposite to the dominant direction), and (3) dots moving in random directions (Figure 1A). The dominant direction was either leftward or rightward, randomized in each trial. The total number of dots in the motion stimuli was fixed to 4,241. Visual stimuli were presented at the center of the screen. Each dot had a diameter of 0.05 degrees and moved at 5 degrees per second. Each dot had a limited lifetime of five frames (83 ms) and was subsequently regenerated at a new location once its lifetime expired. All dots were black and were moving within an invisible circle with a diameter of 6 degrees. A red fixation dot was presented at the center of the screen throughout the experiment.

**Figure 1.**
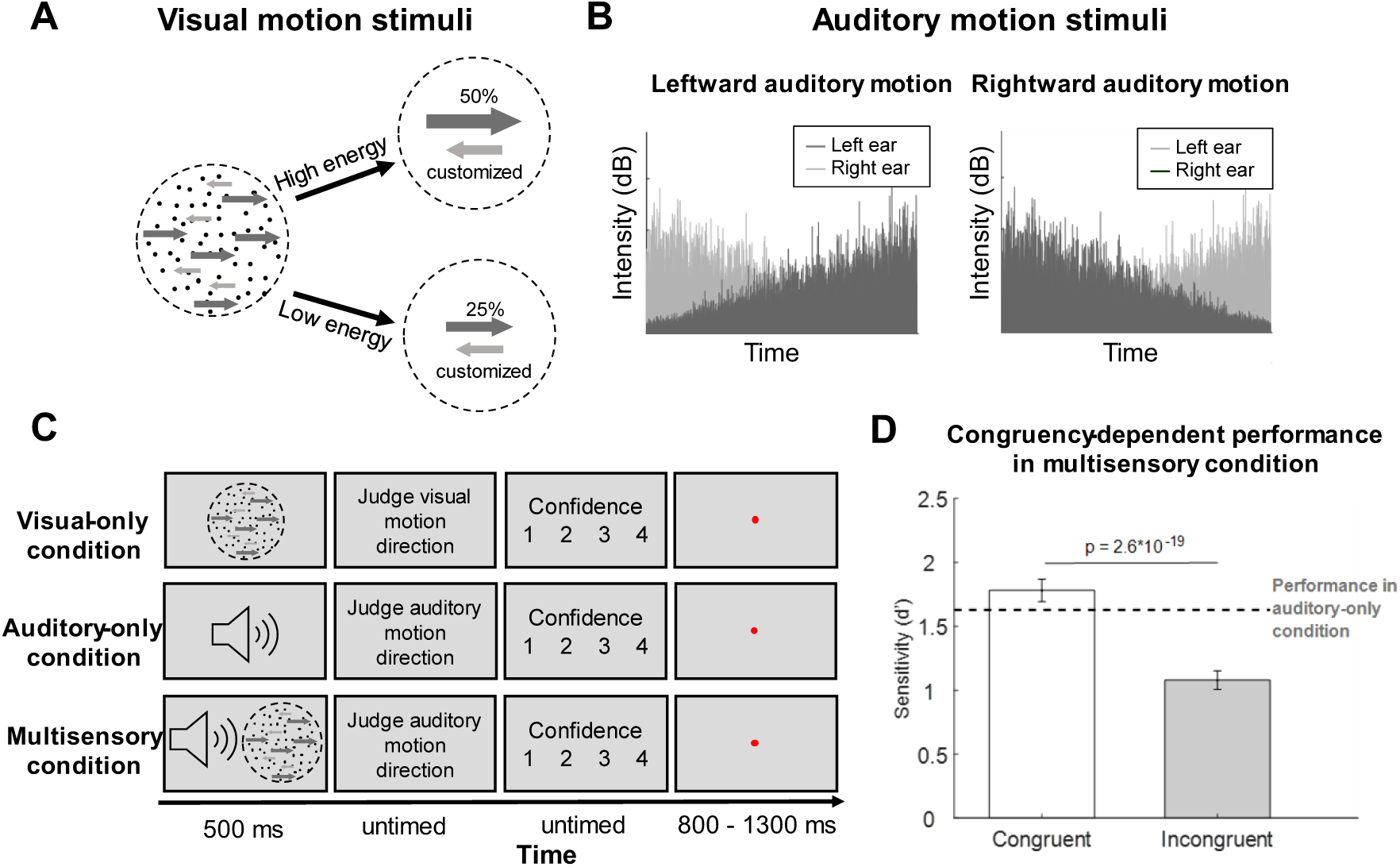
Experimental paradigm and main results. (A) Visual stimuli used in the experiments. The visual stimuli were random-dot kinematograms that consisted of dots moving in a dominant direction (either leftward or rightward for each trial), a non-dominant direction (always opposite to the dominant direction), and random directions. The number of dots moving in the dominant direction was fixed to 50% of the total number of dots for the high-energy stimuli and 25% for the low-energy stimuli. The number of dots moving in the non-dominant direction was customized in different ways for different participants (see Methods). Note that the high coherence in the dominant direction was consistently paired with high coherence in the non-dominant direction. This relationship results in a dissociation between confidence and accuracy. The total number of dots was constant for stimuli with different energy levels. (B) Auditory stimuli used in the experiments. We used cross-faded white noise as auditory motion stimuli. For leftward motion, the sound played to the left ear faded in (i.e., the sound intensity increased over time), while the sound in the right ear faded out (i.e., the sound intensity decreased over time). The opposite was true for rightward motion. (C) Trial structure. Each trial started with motion stimuli (visual-only, auditory-only, or a combination of visual and auditory). Participants then judged the direction of motion (left vs. right) and provided confidence on a 4-point scale. In the multisensory condition, participants judged the auditory motion, but their judgments were typically influenced by the visual motion. The next trial started after a fixation interval of 800-1300 ms. (D) Performance was better for congruent vs. incongruent trials in the multisensory condition.

We created two types of motion stimuli: high-energy stimuli where 50% of dots move in the dominant direction, and low-energy stimuli where 25% of dots move in the dominant direction. Note that the term "energy" here refers to the maximal level of evidence supporting the correct choice. This definition is distinct and unrelated to the concept of "energy" in the Motion Energy model proposed by Adelson & Bergen^38^ To adjust the performance in the high- and low-energy conditions, we customized the percent of dots moving in the non-dominant direction. The experiment included three participant groups to induce different confidence-accuracy dissociations for the visual stimuli. In the first group of participants (N = 24), we aimed to match the performance for high- vs. low-energy stimuli. To do so, we ran 3-down-1-up staircases on the proportion of dots moving in the non-dominant direction separately for the high- and low-energy stimuli. The initial proportion of dots moving in the non-dominant direction was 6% and 3.5% for high- and low-energy stimuli. The initial step sizes of the staircases were 1.5% and 0.9%, and were reduced to 1.0% and 0.6% after three reversals, and to 0.5% and 0.3% after another three reversals. The staircase ended after 150 trials or 14 reversals, whichever came first. The threshold was determined as the average value at the time of the last eight reversals. The final proportions of dots moving in the non-dominant direction were 30.04% (SD = 4.75) for the high-energy condition and 6.98% (SD = 3.18) for the low-energy condition (Figure S1). However, the resulting confidence and accuracy were in the same direction – both confidence and performance were larger for the high- compared to the low-energy stimuli. Therefore, in the second group of participants (N = 25), we aimed to further dissociate confidence and accuracy by inducing lower performance but higher confidence for the high-energy stimuli compared to the low-energy stimuli. For this purpose, we fixed the percent of dots moving in the non-dominant direction to 45% for the high-energy stimuli and 0% for the low-energy stimuli. The manipulation was successful, but the resulting performance for the high-energy condition was quite low. Thus, in the final group of participants (N = 50), we slightly decreased the percent of dots moving in the non-dominant direction to 31% for the high-energy stimuli (and kept it at 0% for the low-energy stimuli). None of the participants were repeated among the three groups. To maximize power, we analyzed the data from all participants together. However, the pattern of results was qualitatively the same for each group when they were analyzed separately (see Figure S1).

The auditory motion stimuli consisted of signal sounds of either leftward or rightward direction and white noise sounds with no direction. To create rightward auditory motion, the intensity of the signal sound in the right ear increased from zero to maximum loudness while the intensity of the signal sound in the left ear decreased from maximum loudness to zero (Figure 1B). The opposite was true for leftward motion. To determine the relative loudness intensity of the signal and noise sounds, we ran a 2-down-1-up staircase on the ratio of the signal sounds to the total loudness (signal + noise sounds), and used the obtained ratio for the target motion in the formal sessions. The starting point of the staircase was 70%; the initial step size was 3%, which was reduced to 2% after three reversals, and to 1% after another three reversals. The staircase ended after 120 trials or 12 reversals, whichever came first. The threshold was determined as the average value at the time of the last six reversals.

### Procedure

The experiment included three different conditions: visual-only, auditory-only, and multisensory. In all cases, participants judged the direction of motion (leftward vs. rightward) and rated their confidence on a 4-point scale (lowest, low, high, highest). The visual-only and auditory-only conditions featured stimuli from a single modality. Critically, the multisensory condition included both visual and auditory stimuli, but participants were instructed to judge the auditory motion direction and ignore the visual motion direction. Nevertheless, based on previous research^29,39–41^, we expected that visual stimuli would affect auditory judgments via automatic integration mechanisms. Motion stimuli were presented for 500 ms and all responses were untimed. After the end of a trial, a fixation circle was shown for a period randomly chosen between 800-1300 ms before the next trial started (Figure 1C).

The experiment began with a screening session intended to remove participants who could not properly perform the visual-only task. During this session, participants completed blocks that contained 25 trials with low-energy stimuli (where we set 0% of dots to move in the non-dominant direction) and 25 trials with high-energy stimuli (where we set 8.3% of dots to move in the non-dominant direction). We passed participants who could perform at 80% correct or better for both types of stimuli. Fourteen participants were unable to reach that threshold for the low-energy stimuli even after up to eight screening blocks and therefore did not participate in the main experiment. These participants are not included in the count of recruited participants. No participants dropped out or declined participation.

The main experiment consisted of two visual-only, one auditory-only, and two multisensory blocks. Each block consisted of 80 trials and the different types of blocks were randomly interleaved for each participant (400 trials total). The visual-only and multisensory blocks contained an equal number of trials with high- and low-energy stimuli. In the multisensory blocks the direction of auditory motion was pseudo-randomized to contain equal number of leftward and rightward motion, whereas the direction of visual motion was fully randomized such that the two stimuli were congruent on average in 50% of the trials. To increase trial number per condition, the third group of participants mentioned above completed four visual-only blocks and 10 blocks where auditory-only and multisensory trials were interleaved (45 trials per block, 630 trials total) but did not need to indicate confidence for the auditory-only and multisensory trials. No one was present during the experiment except the participant and the researcher. The researcher was not blind to the experimental condition and the study hypothesis.

### Apparatus

We presented all visual stimuli on a Dell monitor (47.5 cm x 26.5 cm; refresh rate = 70 Hz) positioned 50 cm away from the participants. We delivered the auditory stimuli through SENNHEISER HD 280 PRO headphones. All sounds had a sampling frequency of 44.1 kHz.

### Behavioral analyses

For each condition, we calculated task performance d’ based on the signal detection theory ^42^ using the formula:

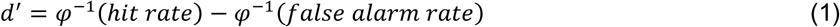

where 𝜑^−1^ denotes to the inverse of the cumulative standard normal distribution converting the hit and false alarm rates to Z scores.

We performed two-sided paired sample t-tests to compare performance d’ and confidence levels between the high- and low-energy stimuli in the visual-only condition, as well as both the visual weights see below for how those were estimated and overall performance d’ in the multisensory condition. Normality and equal variances were formally tested since t-tests were relatively robust to small violations of these assumptions given our large number size (N=99). We report Cohen’s d and the ayes factors^43^ for all t-tests. We calculated Cohen’s d by taking the difference between the mean of the two samples and then dividing it by the pooled standard deviation of the two samples. The default Cauchy priors with a scale parameter of .7071 were used for all Bayes factors analyses. All the statistical analyses were run with MATLAB (MathWorks, Version 2022a).

### Computational model

We developed a model of the computations involved in confidence ratings in the visual-only condition and multisensory integrations in the multisensory condition. The model assumes that both visual and auditory evidence (*x*_*visual*_ and *x*_*auditory*_) are sampled from Gaussian distributions in accordance with signal detection theory^42^ such that leftward and rightward motion stimuli produce distributions of internal evidence coming from 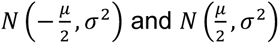, respectively.

Participants give confidence ratings by applying the same criteria for both high- and low-energy stimuli. Critically, the energy level of the visual stimuli could alter the variability of the internal evidence distributions^22,44^. This allows for the high-energy stimuli to have higher variability of the internal signal. In turn, this leads to the high-energy distributions occupying more of the high-confidence regions compared to the low-energy distributions, leading to more high-confidence trials^45–47^ (Figure 3A).

We fit this part of the computational model to the auditory-only and visual-only data in the following manner. For the auditory-only condition, the model had one free parameter for the distance between the peaks of the Gaussian distributions of internal evidence, 𝜇_*auditory*_, with the standard deviation (SD) of the evidence distributions fixed to 1. The parameter 𝜇_*auditory*_ was estimated directly from the data using Equation 1 above. For the visual-only condition, the model had 10 free parameters: the distance between the peaks of the high-energy distributions, 𝜇_𝐻𝐸_, the distance between the peaks of the low-energy distributions, 𝜇_𝐿𝐸_, the SD of the high-energy distributions, 𝜎_𝐻𝐸_, and 7 response criteria 𝑐_𝑖_ with 𝑖 = −3, −2, −1,0,1,2,3. Because the model predictions depend only on the ratio between the SDs for the high- and low-energy stimuli, without loss of generality the SD of the low-energy distributions, 𝜎_𝐿𝐸_, was fixed to 1. The criterion 𝑐_0_ was the decision criterion that separated the two stimulus categories, whereas the rest of the criteria determined the confidence ratings, such that the criteria 𝑐_𝑖_ and 𝑐_−𝑖_ separated the ratings 𝑖 and 𝑖 + 1. Negative criteria separate the confidence ratings for leftward motion decisions, whereas positive criteria separate the confidence ratings for rightward motion decisions. Thus, given the presentation of a stimulus that produces an internal activation 𝑁(𝜇, 𝜎^2^), the probability of confidence rating of 𝑖 for a rightward decision is:

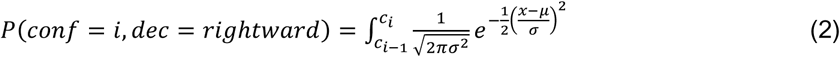

and the probability of confidence rating of 𝑖 for a leftward decision is:

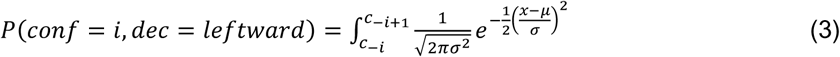

where 𝑐_−4_ = −∞ and 𝑐_4_ = ∞.

So far, we have covered the model’s assumptions regarding the visual-only and auditory-only conditions. For the multisensory condition, we tested three different computations that could potentially underlie multisensory integration.

#### Flexible weight computation

The first computation that could underlie multisensory integration in our task is the Flexible weight computation, according to which participants flexibly combine the visual and auditory signals. The Flexible weight computation includes a free parameter, *w*, that describes the propensity of each participant to use visual information in the multisensory condition. Specifically, the assumption behind this computation is that the auditory signal, *x*_*auditory*_ , is directly combined with the visual signal, *x*_*visual*_ , such that the decision variable for the multisensory decisions, *x*_𝑚𝑢𝑙𝑡𝑖𝑠𝑒𝑛𝑠𝑜𝑟𝑦_ , is:

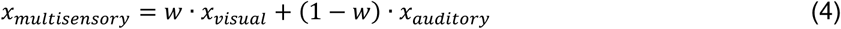

where *w* represents weight allocated to the visual signal, which ranges between 0 and 1 (Figure 3C). Rightward decisions were made for *x*_𝑚𝑢𝑙𝑡𝑖𝑠𝑒𝑛𝑠𝑜𝑟𝑦_ ≥ 0, and leftward decisions for *x*_𝑚𝑢𝑙𝑡𝑖𝑠𝑒𝑛𝑠𝑜𝑟𝑦_ < 0. In an earlier iteration of equation 4, we included a bias term that would allow the Flexible weight computation to account for individual biases for choosing left vs. right motion direction. However, the addition of this term did not improve the overall quality of the model fit or affect any of the model’s critical features. Therefore, to maintain maximum simplicity, the bias term was removed and has not been included for any of the computations underlying multisensory integration.

The Flexible weight computation was implemented using a single free parameter for the weight with which visual stimuli contributed to the multisensory decision variable. Because the distributions for the high-energy visual stimuli extend further to both extremes compared to the distributions for the low-energy visual stimuli, the combination of the visual and auditory signals assumed by the Flexible weight computation results in the high-energy visual stimuli having a stronger impact on auditory judgments.

#### Reliability-weighted computation

Flexible weight computation treats weight as a free parameter, contrasting the approach in Ernst & anks’^2^ seminal paper on cue combination where optimal weight aligns with the reliability of each sensory modality. Unlike Ernst & anks’ study, our participants solely judged auditory motion direction in the multisensory condition, resulting in a zero optimal weight for visual signals. Nevertheless, we tested whether an alternative computation that assumes reliability-weighted averaging of the auditory and visual signals in the multisensory condition describes the data better. We call this Reliability-weighted computation. In the context of our task, the Reliability-weighted computation postulates that the weight (*w*) in equation 4 is not a free parameter but is instead equal to 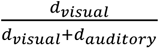, where 𝑑*_visual_* and 𝑑*_auditory_* are the d’ values for the visual and auditory stimuli, respectively. The Reliability-weighted computation thus includes no free parameters and is therefore a lot more constrained than the Flexible weight computation.

#### Flexible causal inference computation

Finally, we considered an alternative computation – which we call Flexible causal inference computation – aimed at mimicking the causal inference mechanism proposed by Körding and colleagues^48^. The idea is that if the auditory and visual signals are sufficiently similar, then they are mandatorily combined based on their reliabilities. However, if they are sufficiently different, then they are judged to have different sources and the participant simply uses the auditory signal (as they should). Because it is not clear a priori how different signals should be before they are judged to have different causes, we implement this computation using a free parameter.

Specifically, the Flexible causal inference computation considers the absolute difference between visual and auditory evidence, 𝑑𝑖𝑓𝑓 = |*x*_*visual*_ − *x*_*auditory*_ |. If 𝑑𝑖𝑓𝑓 > 𝑇 (where 𝑇 is a free parameter representing a threshold), the computation relies on the auditory signal alone, but if 𝑑𝑖𝑓𝑓 ≤ 𝑇, the model uses a mandatory cue combination as in the Reliability-weighted computation. The flexibility in the computation comes from the fact that 𝑇 can take many different values. At the one extreme, 𝑇 can become 0, which means that the visual signal is always disregarded. As the parameter 𝑇 increases, the visual signal is given higher and higher weight in the overall multisensory judgment. At the extreme where 𝑇 takes a very high value (e.g., larger than 6), the difference in the two signals is always less than the threshold and thus the visual signal is always used with a weight of 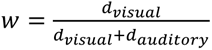. Thus, in practice, the Flexible causal inference computation is very similar to the Flexible weight computation. Indeed, the two computations resulted in relatively similar model fits (Flexible weight computation won by an average of .89 points per participant). More importantly, a model recovery analysis showed that the two computations are not actually distinguishable from each other (Figure S3). Therefore, it appears that the Flexible weight and the Flexible causal inference computations are so strongly related as to be nearly indistinguishable. We therefore present all results related to the Flexible causal inference computation in the Supplementary (Figure S3).

### Model fitting

We performed model fitting using a 2-step procedure. In the first step, we fit the visual-only and auditory-only data, and thus obtained the parameters 𝜇_𝐻𝐸_, 𝜇_𝐿𝐸_, 𝜎_𝐻𝐸_, and 𝜇_*auditory*_. Then, using these parameters, we fit the data from the multisensory condition for each of the three possible computations described above. To fit the model to the data, we used maximum likelihood estimation as in previous studies^22,49,50^. Model fitting was performed using the Bayesian Adaptive Direct Search (BADS) toolbox, version 1.0.5^51^. For both steps, we performed model fitting 10 times and selected the best-fitting iteration as the overall model fit.

### Model comparison

To quantify model fit, we calculated the Akaike Information Criterion (AIC) and Bayesian Information Criterion (BIC). These metrics were computed using the standard formulas: 𝐴𝐼𝐶 = −2 ∗ 𝑙𝑜𝑔𝐿 + 2 ∗ 𝑘 and 𝐵𝐼𝐶 = −2 ∗ 𝑙𝑜𝑔𝐿 + 𝑘 ∗ 𝑙𝑜𝑔(𝑛), where 𝑘 represents the number of free parameters, and 𝑛 is the number of trials. Both AIC and BIC provide a measure of model goodness-of-fit. BIC includes a harsher penalty for model complexity compared to AIC and thus favors simpler models. Lower AIC and BIC values indicate a better fit. To determine whether differences in AIC were statistically significant, we used bootstrapping to generate 95% confidence intervals (CIs) on the summed AIC and BIC differences across models, employing the MATLAB bootci function with 100,000 resamples. Confidence intervals excluding zero suggest a significant difference between models. The model selection based on AIC/BIC is based on a fixed-effects assumption, implying that a single model explains the behavior of all participants.

However, an alternative random-effects approach allows for different models to explain different participants’ behavior. We used the Variational Bayes Analysis toolbox to estimate the model frequencies and their protected exceedance probabilities, which quantify how often a particular model is likely to be the most frequent generative model across the population^52^.

### Model recovery

To assess the distinguishability of the three computations underlying multisensory integration, we performed model recovery analyses. We generated synthetic datasets using each computation, ensuring that the number of participants and trials matched those in the actual experiment. The parameters used for simulating these datasets were derived from the best-fitting parameters.

Each synthetic dataset was then fitted with each of the three computations to determine how well the different computations can be distinguished from each other in our dataset. For each synthetic dataset, we computed AIC and BIC values. To quantify the likelihood that the generating computation is correctly identified, we calculated the frequency with which each computation was selected as the best-fitting computation across all datasets. Specifically, we computed 𝑃_𝑚𝑜𝑑𝑒𝑙_ for every dataset, representing the probability that the computation responsible for generating a particular dataset is correctly identified.

### Parameter recovery

The goal of parameter recovery is to evaluate how accurately the model parameters can be estimated from the data. We utilized the same fits generated during the model recovery analyses but focused only on cases where the same computation was used to generate and later fit the same synthetic dataset. We then compared the estimated parameters to the original known values used in the data generation process. To assess the accuracy of parameter recovery, we calculated the Pearson correlation coefficient between the estimated parameters and the true parameter values across all datasets. A high correlation indicates that the parameters are being accurately recovered by the fitting procedure. Note that the Reliability-weighted computation does not include free parameters; therefore, parameter recovery is not applicable for this computation.

### Weight estimation

As part of our data analysis process, we estimated the weights for the high- and low-energy visual stimuli for each participant in the multisensory condition. We did so for both the empirical data and the data generated by each of the three computations underlying the multisensory integration. To perform this weight estimation, we used a similar procedure as described above for equation 4 except that the weights were separately estimated for high- and low-energy stimuli and the SDs for both high- and low-energy stimulus distributions were set to 1. This procedure determines how much weight is given to each modality when combining signals from different modalities, regardless of the internal variability of each signal.

### Data availability

All data are available at https://osf.io/crkjd/. Data collection for the third group of participants was pre-registered on November 10, 2022. The preregistration can be found at https://osf.io/an7jr.

### Code availability

The experimental codes, codes for analysis, and modelling are available at https://osf.io/crkjd/. All codes were executed using MATLAB (MathWorks, Version 2022a).

## Results

Participants judged the direction of motion (left vs. right) of visual and auditory stimuli. Critically, we used an established method to create two types of visual stimuli that lead to a confidence- accuracy dissociation ^21^. Specifically, “high-energy” visual stimuli had high coherence for both leftward and rightward motion, whereas “low-energy” stimuli had low coherence for both leftward and rightward motion (Figure 1A). Previous research has shown that if one adjusts the motion coherences for leftward and rightward motion appropriately as to match the sensitivity levels d’ for the high- and low-energy stimuli, participants express higher confidence for the high-energy stimuli ^21^.

We had three conditions: visual-only, auditory-only, and multisensory. In the visual-only condition, participants judged the direction of motion of the high- and low-energy visual stimuli described above. In the auditory-only condition, participants experienced auditory stimulation of varying loudness separately in each ear designed to give the impression of leftward or rightward movement and judged the direction of motion (Figure 1B). In the multisensory condition, the visual (both high- and low-energy) and auditory stimuli were presented simultaneously and were congruent 50% of the time. Participants indicated the direction of motion *of the auditory stimuli* irrespective of the direction of visual motion (Figure 1C). In the multisensory condition, the auditory stimuli were randomly paired with either the low- or the high-energy visual stimuli and we tested whether the weight given to the visual stimuli would reflect their corresponding objective reliability or subjective confidence.

We first confirmed that participants integrated the auditory and visual motion stimuli in the multisensory condition. For this, we separately plotted participants’ sensitivity d’ for congruent trials (where visual and auditory directions were the same) and incongruent trials (where visual and auditory directions conflicted) across high- and low-energy visual stimuli. We found that congruent trials produced significantly higher sensitivity than incongruent trials (t(98) = 11.23, p = 2.6*10^-19^, Cohen’s d = 1.13 , 95% CI = [.58, .83], BF_10_ = 2.0*10^16^; Figure 1D). We also observed a small but significant interaction, such that the d’ difference between congruent and incongruent trials was larger for low-energy d’ difference = .80) than for the high-energy trials d’ difference = .61; F(1, 98) = 4.44, p = .04, partial η^2^ = 0.04). In addition, performance was better for congruent trials compared to the auditory-only condition (t(98) = 2.29, p = .02, 95% CI = [.02, .28], Cohen’s d = .23, BF_10_ = 1.34), but worse for incongruent trials compared to the auditory-only condition (t(98) = 8.92, p = 2.6*10^-14^, 95% CI = [.43, .67], Cohen’s d = .90, BF_10_ = 2.7*10^11^). Finally, we also found that reaction times were significantly shorter when visual and auditory motion were congruent compared to incongruent directions (t(197) = 4.74, p = 4.1*10^-6^, 95% CI = [.01, .03], Cohen’s d = .34, BF_10_ = 2875.4; Figure S4). Together, these results show that participants integrated the auditory and visual motion stimuli when making decisions in the multisensory condition instead of the visual stimuli simply interfering with auditory processing.

We then verified that the high- and low-energy visual stimuli produced a confidence-accuracy dissociation. Indeed, in the visual-only condition, high-energy visual stimuli led to lower performance (d’) (t(98) = 5.7, p = 1.0 *10 ^-7^, 95% CI = [.42, .86], Cohen’s d = .58, BF_10_ = 1.2*10^5^, two-sided paired t test; Figure 2A left), but higher confidence (t(98) = 9.1, p = 9.9*10^-15^, 95% CI = [.29, .46], Cohen’s d = .92, BF_10_ = 6.9*10^11^; Figure 2A right) compared to the low-energy visual stimuli.

**Figure 2.**
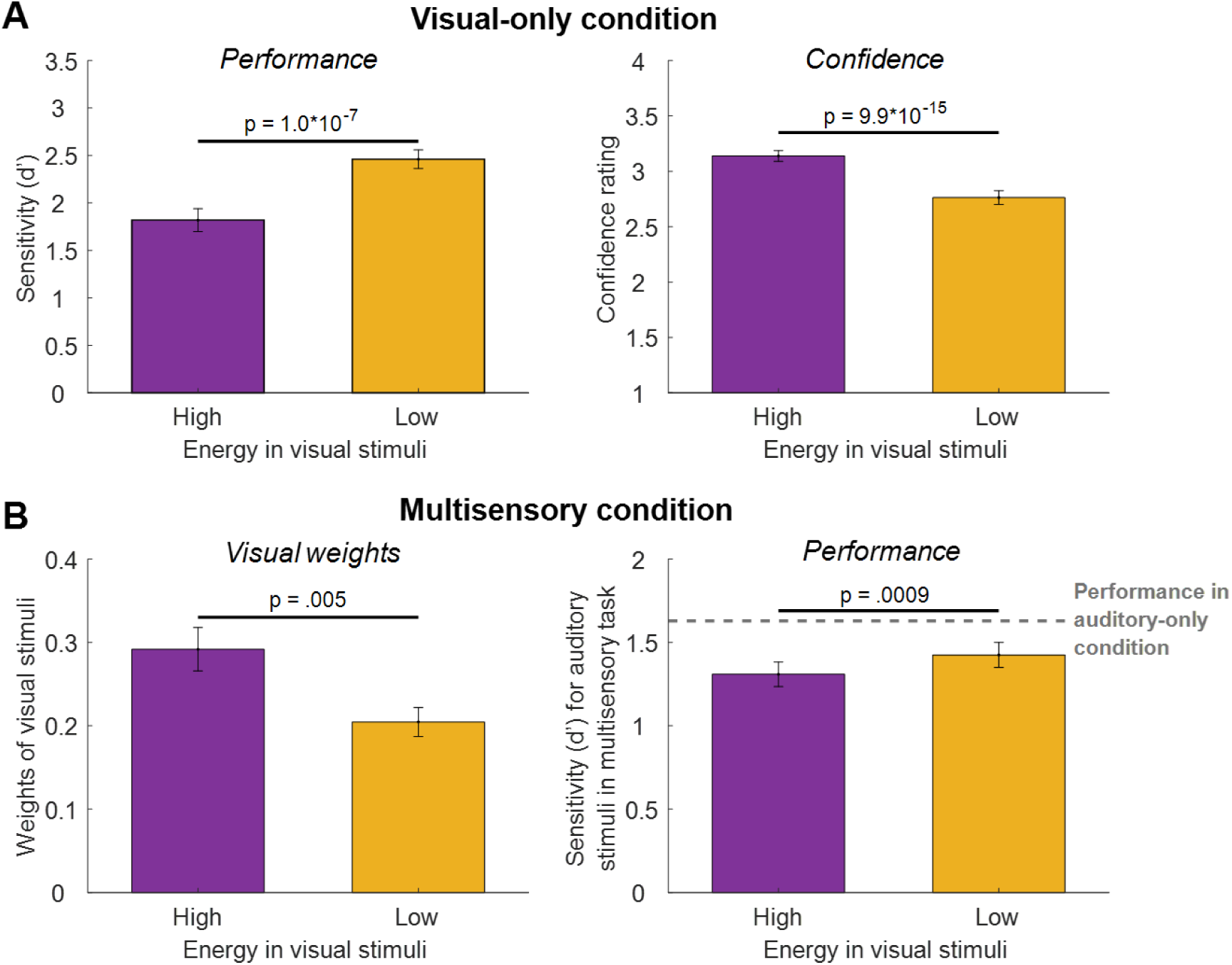
Experimental results. (A) In the visual-only condition, high-energy visual stimuli led to lower performance (left) but higher confidence (right). (B) In the multisensory condition (with congruent and incongruent trials combined), high-energy visual stimuli were weighed more heavily in judgments (left), and both multisensory conditions had lower d’ than the auditory-only condition (dashed line), with a larger decrease for high-energy stimuli (right). Error bars indicate SEM.

Critically, we tested how high- and low-energy visual stimuli affected auditory motion judgment in the multisensory condition. Using the observed performance in the visual-only and auditory-only conditions, we computed the weight of the visual information on the auditory judgments in the multisensory condition. Across all trials, we found that the visual stimuli had a substantial influence (average visual weight = .25; average auditory weight = .75) even though participants were asked to only judge the auditory direction in the multisensory condition. Critically, the weight was substantially higher for the high-energy (average weight = .29) compared to the low-energy stimuli (average weight = .20; t(98) = 3.43, p = .0009, 95% CI = [.04, .14], Cohen’s d = .35, BF_10_ = 25.27; Figure 2B left). Consistent with these results, we also observed that both multisensory conditions exhibited lower d’ than the auditory-only condition (high-energy: t(98) = 5.48, p = 3.4*10^-7^, 95% CI = [.20, .44], Cohen’s d = .55, BF_10_ = 3.9*10^4^; low-energy: t(98) = 3.41, p = .001, 95% CI = [.09, .32], Cohen’s d = .34, BF_10_ = 23.28, Figure 2B right), but that the decrease was larger for the high- compared to the low-energy stimuli (t(98) = 2.90, p = .005, 95% CI = [.04, .20], Cohen’s d = .29, BF_10_ = 5.60). These results demonstrate that multisensory integration was more strongly influenced by the high-energy visual stimuli in line with their higher confidence despite their lower associated performance.

To explain these results, we developed a simple computational model. The model assumes that high-energy visual stimuli produce internal evidence distributions with a significantly larger variance but only a slightly larger distance between the means of the left and right stimulus distributions than low-energy stimuli (Figure 3A). However, as in previous work^45–47^, participants use the same confidence criteria for both stimulus types. This leads to higher confidence ratings for high-energy stimuli despite lower d’ levels, thus explaining the confidence-accuracy dissociation in the visual-only condition. Indeed, the model successfully reproduced the confidence-accuracy dissociation by producing lower performance (t(98) = 2.98, p = .004, 95% CI = [.14, .72], Cohen’s d = .30, BF_01_= 6.92) but higher confidence (t(98) = 6.79, p = 8.8*10^-10^, 95% CI = [.16, .29], Cohen’s d = .68, BF_10_ = 1.1*10^7^) for the high- compared to the low-energy stimuli in the visual-only condition (Figure 4A).

**Figure 3.**
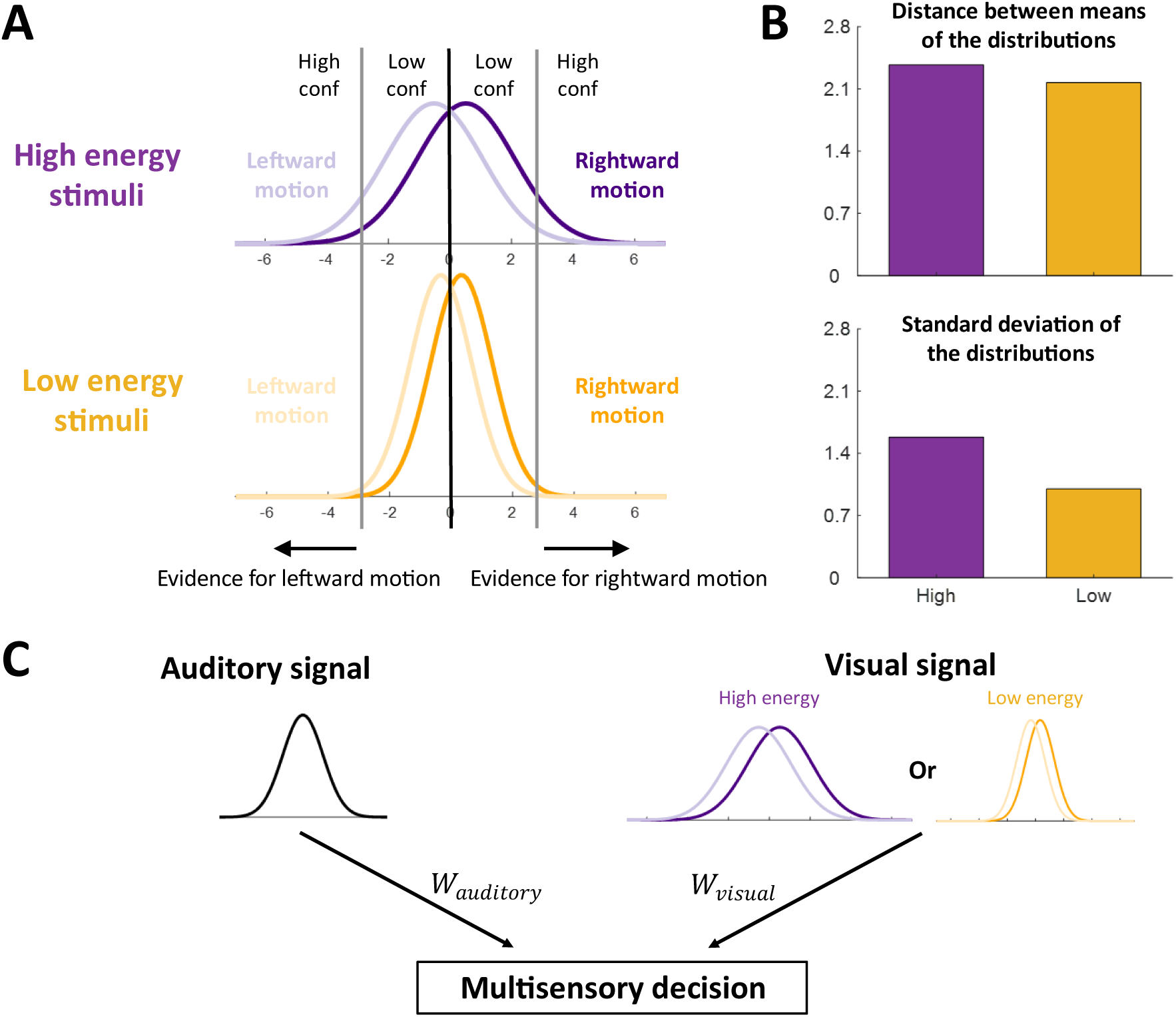
Computational model. (A) Internal distributions of evidence for high- vs. low-energy stimuli. The model assumes that the distribution for high-energy stimuli has a larger variability compared to that of low-energy stimuli, resulting in more trials falling in the range in the tails with higher confidence. The distributions shown in the figure are the average distributions obtained after fitting the model to the data. (B) Standard deviation SD and difference between the two distributions’ means from panel A, showing that high-energy visual stimuli produce internal evidence distributions with a significantly larger variance but only a slightly larger distance between the means of the left and right stimulus distributions than low-energy stimuli. (C) Multisensory-decision model. Visual signals are combined directly with auditory signals without any normalization, such that *x*_*multisensory*_ = *w* · *x*_*visual*_ + (1 − *w*) · *x*_*auditory*_. We tested two main computations underlying multisensory integration. The Flexible weight computation treats the parameter *w* as a free parameter. The Reliability-weighted computation fixes the parameter *w* to the value that would result in weighing each sensory signal according to its reliability^2^.

**Figure 4.**
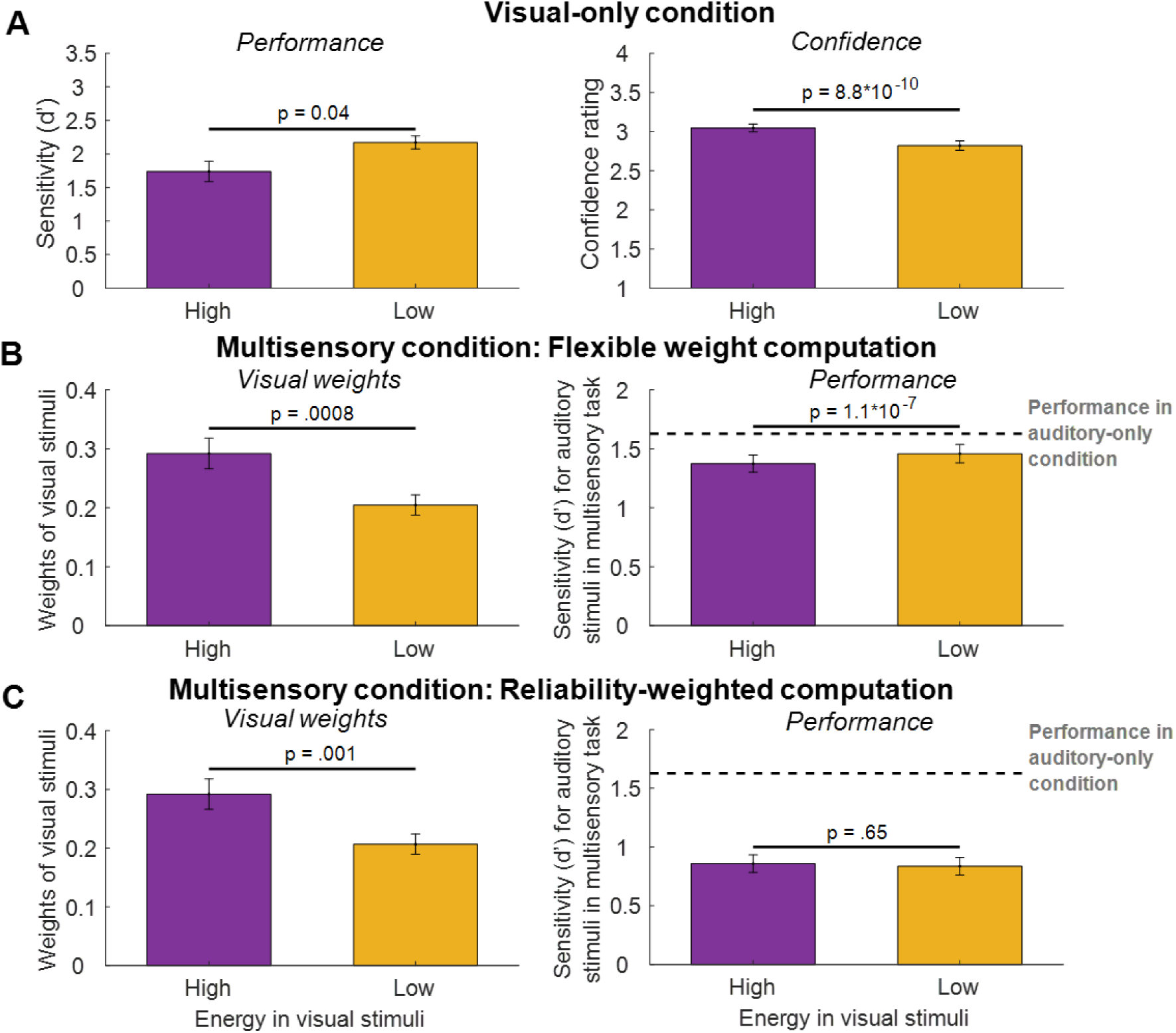
Model fits. (A) he model successfully reproduced lower multisensory d’ for the high-energy stimuli compared to the low-energy stimuli, consistent with the associated higher confidence but lower accuracy for the high-energy stimuli. (B) The Flexible weight computation well reproduced the higher estimated weight and the overall lower multisensory d’ and for the high- compared to the low-energy visual stimuli. (C) The Reliability-weighted computation produced higher weight for the high- compared to the low-energy visual stimuli (left panel). However, it produced similar multisensory d’ for the high- compared to the low-energy visual stimuli. Error bars indicate SEM.

We examined two main computations that can underlie multisensory integration: the Flexible weight computation and the Reliability-weighted computation. The Flexible weight computation posits that participants flexibly combine the visual and auditory signals, with the weight being a free parameter. In contrast, the Reliability-weighted computation assumes that sensory signals are combined based on their respective reliabilities, and the weight is not a free parameter. We fit models instantiating each of these computations to the multisensory data and examined how well each computation explained the observed multisensory effects.

We found that the Flexible weight computation mirrored the behavioral effects, despite assuming a single weight for the low- and high-energy visual stimuli in the multisensory condition.

Specifically, the Flexible weight computation reproduced the higher estimated weight for the high- compared to the low-energy stimuli (t(98) = 3.47, p = .0008, 95% CI = [.04, .14], Cohen’s d = .35, BF_10_ = 28.00) and the overall lower multisensory d’ for the high- compared to the low-energy visual stimuli (t(98) = 5.72, p = 1.1*10^-7^, 95% CI = [.06, .11], Cohen’s d = .58, BF_10_ = 1.1*10^5^; Figure 4B). In contrast, though the Reliability-weighted computation reproduced lower weight for the high- compared to the low-energy visual stimuli (t(98) = 3.36, p = .001, 95% CI = [.04, .14], Cohen’s d = .34, BF_10_ = 20.27), it failed to reproduce the overall lower multisensory d’ for the high- compared to the low-energy visual stimuli (t(98) = .45, p = .65, 95% CI = [-.12, .07], Cohen’s d = .05, BF_01_ = 8.14; Figure 4C). Note that the success of both computations in reproducing the higher visual weights for the high- compared to the low-energy conditions is due to the fact that the high-energy distributions extend further to both extremes (Figure 3A), leading to a stronger influence on the judgments in the multisensory condition. Overall, despite its simplicity, the model with Flexible weight computation was able to explain both the confidence-accuracy dissociation and the multisensory integration bias by postulating a unified computational principle.

In close correspondence to its better qualitative fits (Figure 4B,C), the Flexible weight computation outperformed the Reliability-weighted computation by a total of 8,852 BIC points (Figure 5A; see Figure S5 for AIC results, which show an even bigger advantage for the Flexible weight computation). The Flexible weight computation also exhibited high parameter recoverability (r = 0.92, p = 1.0 × 10^-42^, 95% CI = [.88, .94], Figure 5B). We also observed high model recovery, such that the correct computation was recovered an average of 90.41% (Figure 5C; see Figure S5 for AIC results).

**Figure 5.**
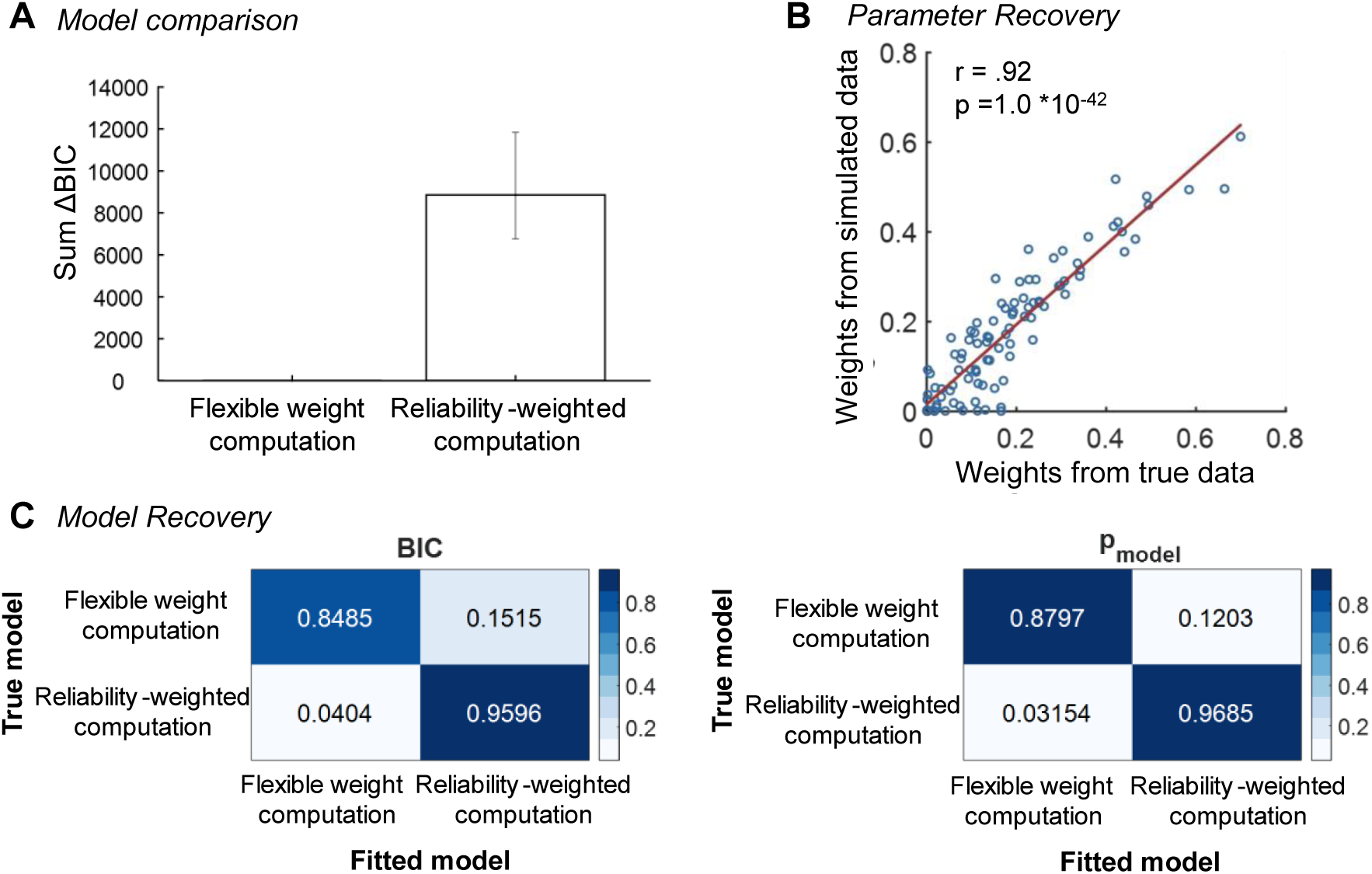
Model comparison, parameter recovery, and model recovery. (A) Comparison of model performance between the Flexible weight computation and the Reliability-weighted computation based on BIC values. The Flexible weight computation outperformed the Reliability-weighted computation. Error bars indicate 95% confidence intervals (CIs) generated using bootstrapping. (B) Parameter recovery for the weight parameter (*w*) of the Flexible weight computation. Pearson’s correlation between weights fitted from simulated data and true data demonstrates effective parameter recovery. The red line represents the fit using a linear regression model. (C) Model recovery analysis for the Flexible weight and Reliability-weighted computations. Model recovery was assessed using standard fixed-effects analyses (left) and using random-effects modeling (right). In both cases, we observe excellent model recovery showing that the two computations are clearly distinguishable from each other.

Finally, we also tested whether the multisensory results can be explained by a Flexible causal inference computation. This computation first determines if the visual and auditory signals likely originate from the same or different sources based on the absolute difference between these signals (a free parameter). When the signals are judged to have different sources, the computation relies solely on the auditory signal, whereas when the signals are judged to have the same source, the two signals are combined based on their reliabilities. However, we found that the Flexible causal inference computation strongly mimicked the Flexible weight computation.

Specifically, the two computations were easily confusable, showing very poor model recovery (Figure S3). This is likely due to the fact that their free parameters allow them to behave similarly for the majority of the range of these free parameters (see Methods).

## Discussion

Using an established manipulation for producing confidence-accuracy dissociations^21^, we found that multisensory integration follows subjective confidence instead of objective performance^5,53,54^. Critically, by using stimuli that specifically dissociate sensitivity and confidence, we show that the weight given to visual stimuli in multisensory trials follows the subjective confidence ratings instead of the objective sensitivity associated with these stimuli. We note that in most cases, objective uncertainty and subjective confidence in standard 2-choice tasks go together, such that confidence is typically higher for conditions with higher accuracy^13,14,55^. However, in the cases where confidence and accuracy do dissociate (e.g., for a condition that produces higher confidence despite lower accuracy), then it is the confidence that drives multisensory integration.

Our work suggests the existence of common computations underlying multisensory integration and metacognitive confidence reports. Several previous papers have also examined whether metacognitive confidence judgments share computations with other processes. For example, recent work has demonstrated the existence of similar computational noise across cognition, metacognition, and even meta-metacognition^56,57^ as well as similar neural correlates for perceptual decision making and confidence^58^. In addition, the metacognitive ability to provide confidence ratings predictive of one’s accuracy is at least partly domain-general^59,60^. Importantly, none of this previous work has examined whether automatic sensory inference performed within sensory areas of the brain may also share mechanisms with the deliberate computations associated with higher-order cognition. The current findings thus provide critical evidence for the existence of common computations among divergent process related to perception.

Our computational model implies that internal evidence distributions with higher variance lead to both higher confidence and higher weighing in multisensory judgments. At first, this may appear counterintuitive because in traditional cue combination studies higher variance means lower performance. The difference stems from the fact that we used a 2-choice categorization task rather than an estimation task. In estimation tasks, higher variability occurs for more uncertain stimuli, which leads to lower confidence^16,65^ and less weight in multisensory judgments^2,6^.

However, in 2-choice tasks, the uncertainty is jointly determined by the variance of the internal distributions and the distance between their means^66^. Thus, in a 2-choice task, conditions with high variance of the internal distributions can be associated with high performance as long as the higher variability is offset by a larger distance between the means of the distributions (as is the case in our model; Figure 2A, B).

The multisensory results observed here were well fit by assuming a flexible combination of the visual and auditory stimuli (Flexible weight computation). Notably, very similar results were obtained by assuming that visual and auditory stimuli are flexibly combined using the principles of causal inference (Flexible causal inference computation)^48^. Indeed, our model recovery analyses demonstrated that these two computations are almost indistinguishable in the context of the current experiment. The underlying reason for the success of both the Flexible weight and Flexible causal inference computations is that both are primarily influenced by the differing variances of high vs. low-energy visual-only stimuli, where visual motion stimuli with greater variance exert a larger impact on auditory motion judgments. Therefore, the conclusion that multisensory integration follows subjective confidence rather than objective performance is supported by both of these potential computations underlying multisensory judgments.

The finding that multisensory integration is driven by confidence supports the hypothesis of a late cortical origin for multisensory weighting processes^67^. Previous studies provide evidence for the involvement of both sensory and parieto-frontal regions in multisensory integration^68–72^. To further explore the neural signatures of how confidence influences this process, future research could examine differences in neural responses to high- versus low-energy stimuli during multisensory integration using techniques such as EEG and fMRI.

We used a specific paradigm that induces a confidence-accuracy dissociation^21^ because this paradigm can be adapted to a multisensory study. Many other manipulations for inducing confidence-accuracy dissociations have been developed in the literature^22,73–75^ but they are harder to adapt to sensory cue integration. Moreover, here we focus on participants’ propensity to misestimate confidence instead of the noise inherent in providing confidence judgments^49,74,76,77^. Future work should replicate our results using other paradigms that can induce confidence-accuracy dissociation and examine the influence on metacognitive noise.

In conclusion, our work demonstrates that subjective confidence, not objective performance, guides multisensory integration. Our results suggest the existence of common computational mechanisms across vastly different stages of perceptual decision making and may point to the existence of unified inference mechanisms throughout the cortex.

### Limitations

The task used here differs from Ernst & anks’^2^ seminal paper on cue combination, as well as many other multisensory cue combination studies^1,4^. In our work, the task in the multisensory trials was to only judge the auditory signal and ignore the visual stimulus. Optimal performance would therefore be obtained if the visual stimuli were completely ignored (i.e., given weight of 0). It is therefore not surprising that the Reliability-weighted computation – which was developed as a model for cases where optimal performance is achieved via reliability-weighted cue combination – did not fit well with our data. It is an open question whether our conclusions apply to traditional multisensory cue combination that uses estimation tasks. Based on the current results, we predict that subjective confidence would also play a crucial role in multisensory cue combination.

Specifically, if a modality is associated with low reliability but high subjective confidence, it is likely that this modality will be overweighed (relative to optimal) in cue combination studies. Future work should test this prediction, as well as whether this type of effect may explain previous findings of suboptimality in cue combination studies^3,53,54,61–64^.

## Acknowledgments

We thank Minzhi Wang for his help with data collection. This work was supported by the National Institute of Health (award: R01MH119189) and the Office of Naval Research (award: N00014-20-1-2622).

## Author Contributions

Y.G. and D.R. designed research; Y.G. performed research; Y.G. and K.X. analyzed data; and Y.G., D.R., and B.O. wrote the paper.

## Supplementary Results

We employed a linear mixed-effects model to assess multisensory effects across different participant groups. The analysis revealed no statistically significant interaction between the group and condition (F(2,192) = 0.60, p = 0.55, partial 𝜂^2^ = .0062). The main effect of the group was significant (F(2,192) = 8.41,p = 0.0003, partial 𝜂^2^ = .08), indicating significant differences in multisensory effect among the three groups. However, there was no statistically significant main effect of condition (F(1,192) = 0.69, p = 0.408, partial 𝜂^2^ = .004). These results suggest that while the groups differ in their multisensory effect, these differences are consistent across both high and low-energy conditions. Post-hoc comparisons indicate that there are significant differences in multisensory effect between Group 1 and Group 2 (F(1,192) = 8.92, p = 0.0032, partial 𝜂^2^ = .04) and between Group 2 and Group 3 (F(1,192)=16.23, p = 0.08, partial 𝜂^2^ = .08), but not between Group 1 and Group 3 (F(1,192) = 0.36, p = 0.5495, partial 𝜂^2^ = .002). Given that the group differences were consistent across conditions, we opted to combine the groups in the main analysis.

## Supplementary Figures

**Figure S1.**
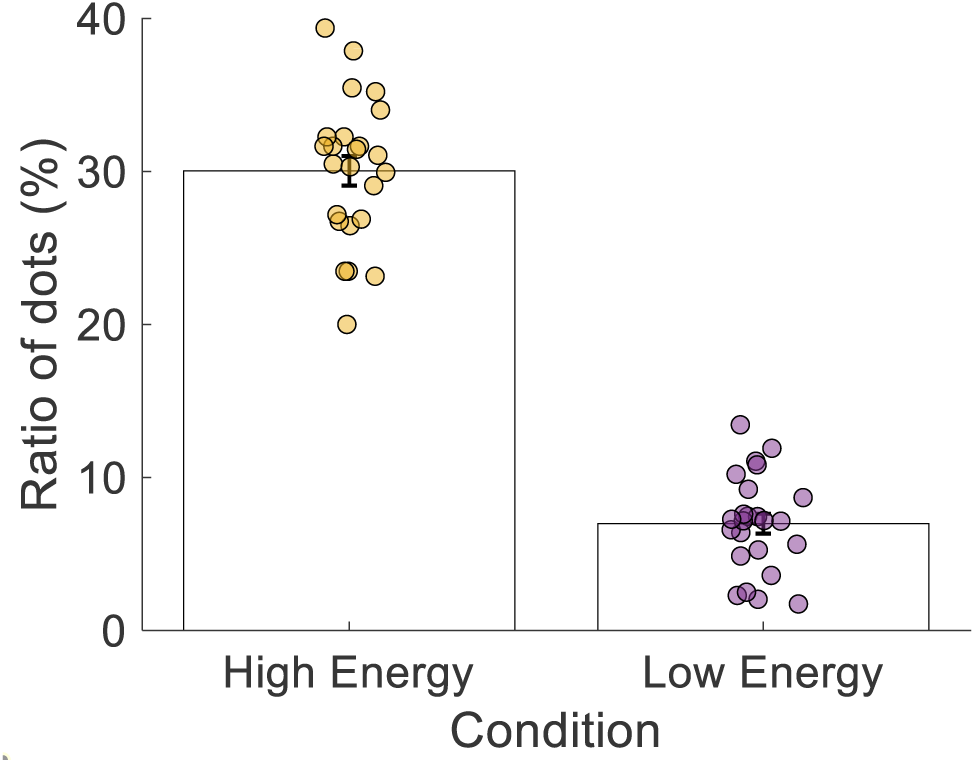
Distributions of percentages of dots moving in the non-dominant direction for high- vs. low-energy conditions across participants in Group 1. In the high-energy condition, the mean percentage of dots moving in the non-dominant direction was 30.04% (SD = 4.75), while in the low-energy condition, the mean percentage was 6.98% (SD = 3.18). For Group 2, the percentage of dots moving in the non-dominant direction was fixed at 45% for high-energy stimuli and 0% for low-energy stimuli. For Group 3, the percentage of dots in the non-dominant direction was slightly decreased to 31% for high-energy stimuli and again set to 0% for low-energy stimuli. Error bars represent the standard error of the mean (SEM).

**Figure S2.**
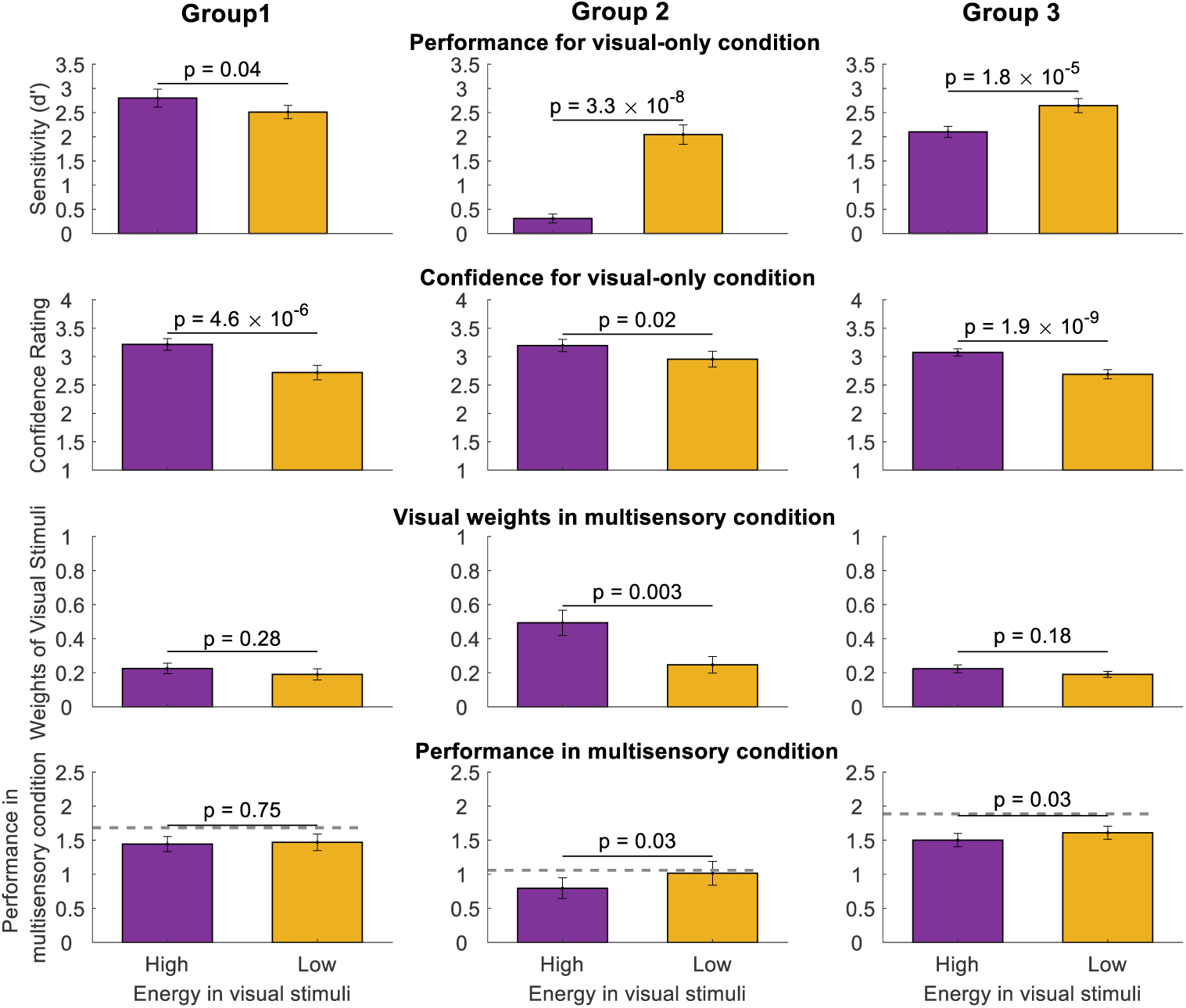
Results for the three participant groups. In the first group of participants (left column), we attempted to match the performance for high- vs. low-energy stimuli by running a 3-down-1-up staircases on the proportion of dots moving in the non-dominant direction separately for the high- vs. low-energy stimuli. The staircase worked reasonably well, but performance was nonetheless slightly but significantly higher for the high-energy visual stimuli (t(23) = 2.18, p = .04, 95% CI [.01, .56], Cohen’s d = .44; top row). Nevertheless, confidence was much higher for the high-energy visual stimuli (t(23) = 5.95, p = 4.6*10^-6^, 95% CI [.32, 67], Cohen’s d = 1.21; second row), similar to what we observed in the pooled data in the main paper. The directions of the visual weights and performance in multisensory condition are the same as in the pooled data but not significant in this group alone. Specifically, the visual stimuli were weighted numerically higher for the high- compared to the low-energy visual stimuli, but this effect was not statistically significant (t(23) = 1.10, p = .28, 95% CI [-.03, 10], Cohen’s d = .22 ; third row). Similarly, the performance in the multisensory condition was lower for the high- compared to the low-energy stimuli, but this effect was also not statistically significant (t(23) = .32, p = .75, 95% CI [-.19, .14], Cohen’s d = .07; last row). In the second group of participants (middle column), we fixed the percent of dots moving in the non-dominant direction to 45% for the high-energy stimuli and 0% for the low-energy stimuli. All results in this group are equivalent to the results in the main text where we pooled data from all three groups. High-energy visual stimuli led to lower performance (t(24) = 7.98, p = 3.3*10^-8^, 95% CI [1.29, 2.18], Cohen’s d = 1.60 but higher confidence (t(24) = 2.61, p = .02,95% CI [.05, 43], Cohen’s d =.52). The visual stimuli were weighted more heavily in multisensory judgments for the high- compared to the low-energy visual stimuli (t(24) = 3.31, p = .003, 95% CI [.09, 40], Cohen’s d = .66. The performance in the multisensory condition was lower for the high- compared to the low-energy stimuli (t(24) = 2.25, p = .03, 95% CI [.02, 42], Cohen’s d =.45). In the third group of participants (right column), we fixed the percent of dots moving in the non-dominant direction to 31% for the high-energy stimuli and 0% for the low-energy stimuli. Results for this group are equivalent to the results in the main text where we pooled the data from all three groups, except that the difference in visual weights used in the multisensory condition for high- vs. low-energy stimuli did not reach significance. High-energy visual stimuli led to lower performance (t(49) = 4.76, p = 1.8*10^-5^, 95% CI [.31, 77], Cohen’s d = .67) but higher confidence (t(49) = 7.35, p = 1.9*10^-9^, 95% CI [.28, 49], Cohen’s d = 1.04) The visual stimuli were weighted numerically higher for the high- compared to the low-energy visual stimuli, but this effect was not statistically significant (t(49) = 1.37, p = .18, 95% CI [-.02, 08], Cohen’s d = .19 . he performance in the multisensory condition was statistically significantly lower for the high- compared to the low-energy stimuli (t(49) = 2.21, p = .03, 95% CI [.01, .21], Cohen’s d =.31). Error bars indicate SEM, dashed line indicates average performance in the auditory-only condition.

**Figure S3.**
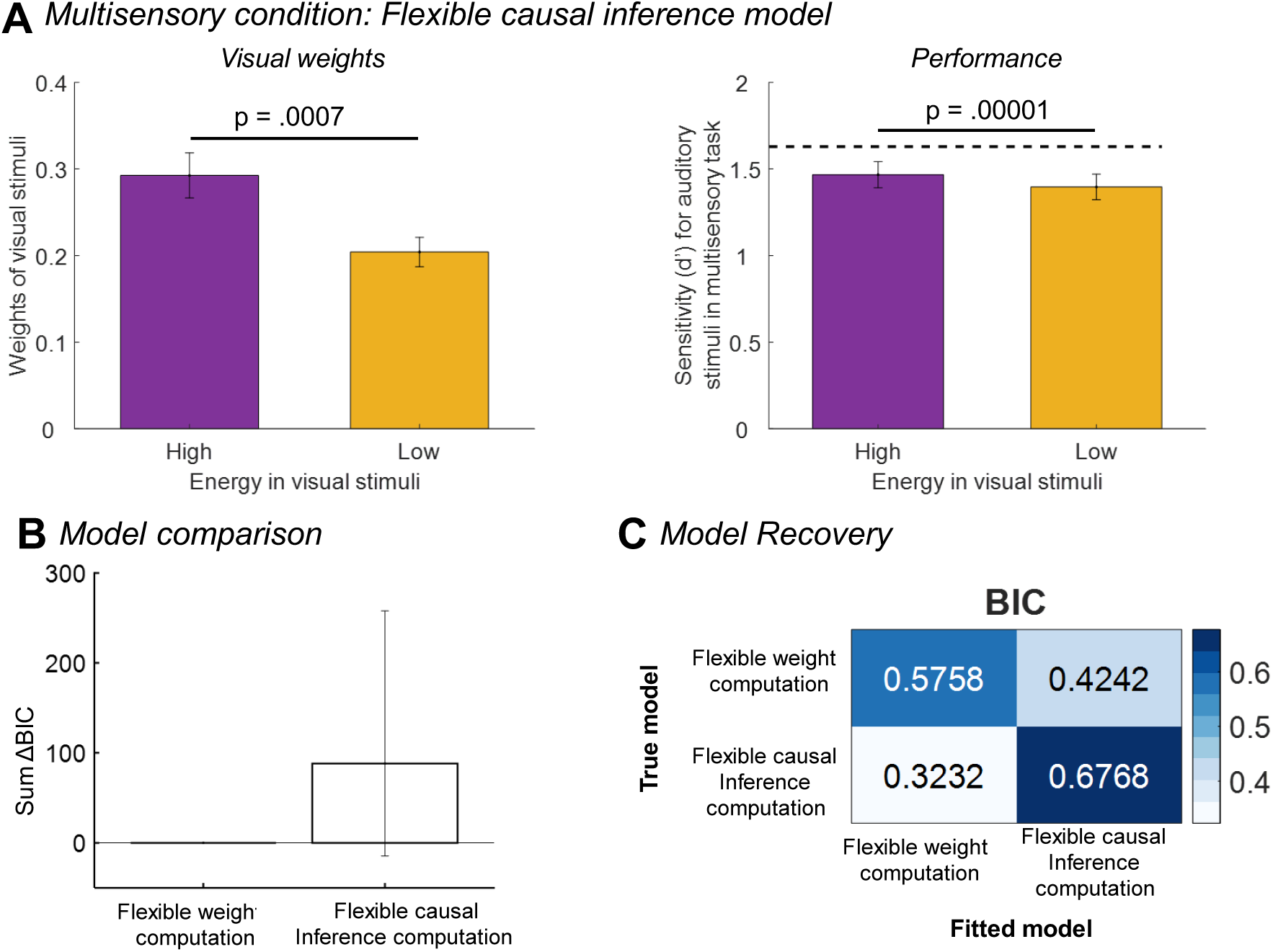
Results for the Flexible causal inference computation. The Flexible causal inference computation postulates that multisensory judgments are made by considering the absolute difference between visual and auditory evidence. If this difference exceeds a certain threshold *T* (a free parameter representing the point at which the cues are considered too dissimilar), the model relies exclusively on the auditory signal. However, when the absolute difference is less than or equal to *T*, the model combines both visual and auditory signals using a reliability-weighted approach. Specifically, the weight assigned to the visual cue is calculated as 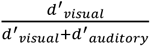, where d′ values represent the sensitivity of each sensory modality. (A) The Flexible causal inference computation reproduced the higher estimated weight for the high- compared to the low-energy stimuli (t(98) = 3.51, p = .0007, 95% CI [-0.03, -.01], Cohen’s d = .35, BF_10_ = 31.26). However, in contrast to the data, the model produced larger multisensory d’ for the high- compared to the low-energy visual stimuli (t(98) = 4.61, p = .00001, 95% CI [-0.03, -.01], Cohen’s d = .46, BF_10_ = 1339.6). (B) Comparison of model performance between the Flexible weight computation and the Flexible causal inference computation based on BIC values. The two computations provided relatively similar model fits. The Flexible weight computation won by a total of 88.2 points, which is an average of .89 points per participant). Error bars indicate 95% confidence intervals (CIs) generated using bootstrapping. (C) Model recovery analysis comparing the Flexible weight and Flexible causal inference computations. The model recovery analysis showed that the two models are not easily distinguishable from each other, with an average model recovery rate of 62.63%. Note that because the Flexible weight and Flexible causal inference computations have the same number of free parameters (one for each), using AIC instead of BIC in panels B and C produces the same results.

**Figure S4.**
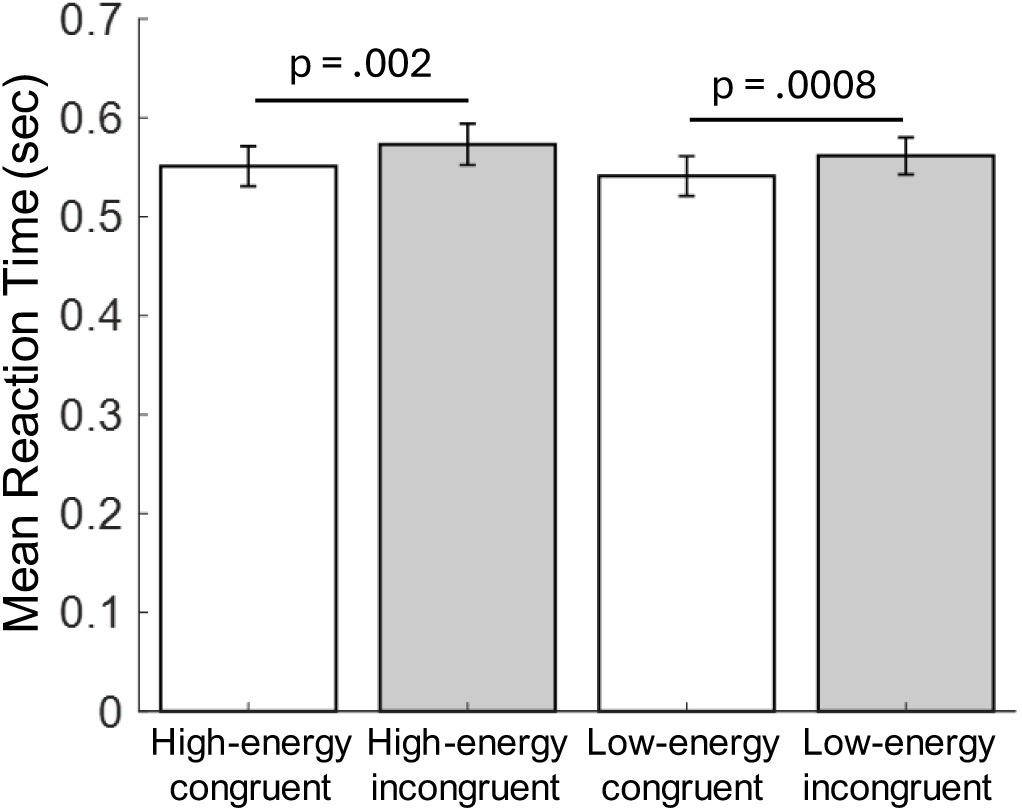
Reaction times for congruent vs. incongruent trials. The reaction times were shorter when visual motion direction and auditory motion direction were congruent compared to incongruent. This effect was significant for both high- and low-energy conditions (High-energy: t(98) = 3.26, p = . 002, 95% CI [.009, .036], Cohen’s d = .33; Low-energy: t(98) = 3.46, p = .0008, 95% CI [.009, .032], Cohen’s d = .35) Error bars indicate SEM.

**Figure S5.**
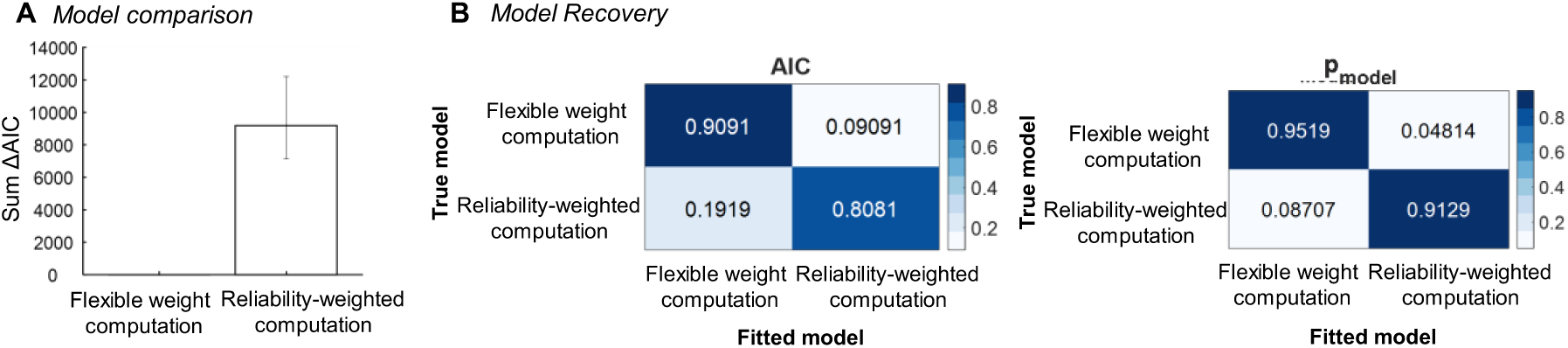
Model comparison and model recovery based on AIC between the Flexible weight computation and the Reliability-weighted computation. (A) Comparison of model performance between the Flexible weight computation and the Reliability-weighted computation based on AIC values. The Flexible weight computation outperformed the Reliability-weighted computation by a total of 9188 AIC points. Error bars indicate 95% confidence intervals (CIs) generated using bootstrapping. (B) Model recovery analysis comparing the Flexible weight and Reliability-weighted computations. The model recovery analysis shows a slight bias towards recovering the Flexible weight computation more frequently. Consequently, to avoid this bias, we used BIC in the main text (BIC showed a slight bias towards the Reliability-weighted computation).

## References

1. Alais, D. & Burr, D. The Ventriloquist Effect Results from Near-Optimal Bimodal Integration. Current Biology 14, 257–262 (2004).

2. Ernst, M. O. & Banks, M. S. Humans integrate visual and haptic information in a statistically optimal fashion. Nature 415, 429–433 (2002).

3. Fetsch, C. R., Pouget, A., Deangelis, G. C. & Angelaki, D. E. Neural correlates of reliability-based cue weighting during multisensory integration. Nat Neurosci 15, 146–154 (2012).

4. Kim, R., Peters, M. A. K. & Shams, L. 0 + 1 > 1: How Adding Noninformative Sound Improves Performance on a Visual Task. Psychol Sci 23, 6–12 (2012).

5. Landy, M. S., Banks, M. S. & Knill, D. C. Ideal-Observer Models of Cue Integration. in Sensory Cue Integration 5–29 (Oxford University Press, 2011). doi:10.1093/acprof:oso/9780195387247.003.0001.

6. Morgan, M. L., DeAngelis, G. C. & Angelaki, D. E. Multisensory Integration in Macaque Visual Cortex Depends on Cue Reliability. Neuron 59, 662–673 (2008).

7. Alink, A. et al. Auditory Motion Capturing Ambiguous Visual Motion. Front Psychol 2, (2012).

8. Wuerger, S. M., Hofbauer, M. & Meyer, G. F. The integration of auditory and visual motion signals at threshold. Percept Psychophys 65, 1188–1196 (2003).

9. Meyer, G. F. & Wuerger, S. M. Cross-modal integration of auditory and visual motion signals. Neuroreport 12, 2557–2560 (2001).

10. Soto-Faraco, S., Spence, C. & Kingstone, A. Cross-Modal Dynamic Capture: Congruency Effects in the Perception of Motion Across Sensory Modalities. J Exp Psychol Hum Percept Perform 30, 330–345 (2004).

11. Spence, C. & Walton, M. On the inability to ignore touch when responding to vision in the crossmodal congruency task. Acta Psychol (Amst) 118, 47–70 (2005).

12. Fleming, S. M., Dolan, R. J. & Frith, C. D. Metacognition: computation, biology and function. Philosophical Transactions of the Royal Society B: Biological Sciences 367, 1280– 1286 (2012).

13. Mamassian, P. Visual Confidence. Annu Rev Vis Sci **2**, 459–481 (2016).

14. Rahnev, D. Visual metacognition: Measures, models, and neural correlates. American Psychologist 76, 1445–1453 (2021).

15. Yeung, N. & Summerfield, C. Metacognition in human decision-making: confidence and error monitoring. Philosophical Transactions of the Royal Society B: Biological Sciences 367, 1310–1321 (2012).

16. Geurts, L. S., Cooke, J. R. H., van Bergen, R. S. & Jehee, J. F. M. Subjective confidence reflects representation of Bayesian probability in cortex. Nat Hum Behav 6, 294–305 (2022).

17. Sanders, J. I., Hangya, B. & Kepecs, A. Signatures of a Statistical Computation in the Human Sense of Confidence. Neuron 90, 499–506 (2016).

18. Maniscalco, B. & Lau, H. A signal detection theoretic approach for estimating metacognitive sensitivity from confidence ratings. Conscious Cogn 21, 422–430 (2012).

19. Mueller, S. T. & Weidemann, Christoph. T. Decision noise: An explanation for observed violations of signal detection theory. Psychon Bull Rev 15, 465–494 (2008).

20. Shekhar, M. & Rahnev, D. Sources of Metacognitive Inefficiency. Trends Cogn Sci 25, 12– 23 (2021).

21. Odegaard, B. et al. Superior colliculus neuronal ensemble activity signals optimal rather than subjective confidence. Proc Natl Acad Sci U S A 115, E1588–E1597 (2018).

22. Rahnev, D. A robust confidence–accuracy dissociation via criterion attraction. Neurosci Conscious 2021, (2021).

23. Samaha, J., Barrett, J. J., Sheldon, A. D., LaRocque, J. J. & Postle, B. R. Dissociating Perceptual Confidence from Discrimination Accuracy Reveals No Influence of Metacognitive Awareness on Working Memory. Front Psychol 7, (2016).

24. Vlassova, A., Donkin, C. & Pearson, J. Unconscious information changes decision accuracy but not confidence. Proceedings of the National Academy of Sciences 111, 16214–16218 (2014).

25. Zylberberg, A., Barttfeld, P. & Sigman, M. The construction of confidence in a perceptual decision. Front Integr Neurosci 6, (2012).

26. Van der Burg, E., Olivers, C. N. L., Bronkhorst, A. W. & Theeuwes, J. Pip and pop: Nonspatial auditory signals improve spatial visual search. J Exp Psychol Hum Percept Perform 34, 1053–1065 (2008).

27. Bertelson, P., Vroomen, J., De Gelder, B. & Driver, J. The ventriloquist effect does not depend on the direction of deliberate visual attention. Percept Psychophys 62, 321–332 (2000).

28. Helbig, H. B. & Ernst, M. O. Visual-haptic cue weighting is independent of modality- specific attention. J Vis 8, 21 (2008).

29. Soto-Faraco, S., Spence, C. & Kingstone, A. Assessing automaticity in the audiovisual integration of motion. Acta Psychol (Amst) 118, 71–92 (2005).

30. Talsma, D., Senkowski, D., Soto-Faraco, S. & Woldorff, M. G. The multifaceted interplay between attention and multisensory integration. Trends Cogn Sci 14, 400–410 (2010).

31. Nidiffer, A. R., Stevenson, R. A., Krueger Fister, J., Barnett, Z. P. & Wallace, M. T. Interactions between space and effectiveness in human multisensory performance. Neuropsychologia 88, 83–91 (2016).

32. Harrar, V., Harris, L. R. & Spence, C. Multisensory integration is independent of perceived simultaneity. Exp Brain Res 235, 763–775 (2017).

33. Juan, C. et al. The variability of multisensory processes of natural stimuli in human and non-human primates in a detection task. PLoS One 12, e0172480 (2017).

34. Rausch, M., Zehetleitner, M., Steinhauser, M. & Maier, M. E. Cognitive modelling reveals distinct electrophysiological markers of decision confidence and error monitoring. Neuroimage 218, 116963 (2020).

35. Rausch, M., Hellmann, S. & Zehetleitner, M. Confidence in masked orientation judgments is informed by both evidence and visibility. Atten Percept Psychophys 80, 134–154 (2018).

36. Xue, K., Zheng, Y., Rafiei, F. & Rahnev, D. The timing of confidence computations in human prefrontal cortex. Cortex 168, 167–175 (2023).

37. Shekhar, M. & Rahnev, D. Distinguishing the Roles of Dorsolateral and Anterior PFC in Visual Metacognition. The Journal of Neuroscience 38, 5078–5087 (2018).

38. Adelson, E. H. & Bergen, J. R. Spatiotemporal energy models for the perception of motion. Journal of the Optical Society of America A 2, 284 (1985).

39. Alink, A. et al. Auditory motion capturing ambiguous visual motion. Front Psychol 3, (2012).

40. Kitagawa, N. & Ichihara, S. Hearing visual motion in depth. Nature 416, 172–174 (2002).

41. Soto-Faraco, S., Lyons, J., Gazzaniga, M., Spence, C. & Kingstone, A. The ventriloquist in motion: Illusory capture of dynamic information across sensory modalities. Cognitive Brain Research 14, 139–146 (2002).

42. Green, D. M. & Swets, J. A. Signal Detection Theory and Psychophysics. (John Wiley, 1966).

43. 43. Krekelberg, B. BayesFactor. Preprint at 10.5281/zenodo.7006300 (2022).

44. Shekhar, M. & Rahnev, D. Human-like dissociations between confidence and accuracy in convolutional neural networks. PLoS Comput Biol 20, e1012578 (2024).

45. Rahnev, D. et al. Continuous theta burst transcranial magnetic stimulation reduces resting state connectivity between visual areas. J Neurophysiol 110, 1811–1821 (2013).

46. Rahnev, D. et al. Attention induces conservative subjective biases in visual perception. Nat Neurosci 14, 1513–1515 (2011).

47. Rahnev, D., Bahdo, L., De Lange, F. P. & Lau, H. Prestimulus hemodynamic activity in dorsal attention network is negatively associated with decision confidence in visual perception. J Neurophysiol 108, 1529–1536 (2012).

48. Körding, K. P. et al. Causal Inference in Multisensory Perception. PLoS One 2, e943 (2007).

49. Shekhar, M. & Rahnev, D. The nature of metacognitive inefficiency in perceptual decision making. Psychol Rev 128, 45–70 (2021).

50. Yeon, J. & Rahnev, D. The suboptimality of perceptual decision making with multiple alternatives. Nat Commun 11, (2020).

51. Acerbi, L. & Ma, W. J. Practical Bayesian Optimization for Model Fitting with Bayesian Adaptive Direct Search. Advances in Neural Information Processing Systems 30, 1834– 1844 (2017).

52. Daunizeau, J., Adam, V. & Rigoux, L. VBA: A Probabilistic Treatment of Nonlinear Models for Neurobiological and Behavioural Data. PLoS Comput Biol 10, e1003441 (2014).

53. Arnold, D. H., Petrie, K., Murray, C. & Johnston, A. Suboptimal human multisensory cue combination. Sci Rep 9, 5155 (2019).

54. Knill, D. C. & Saunders, J. A. Do humans optimally integrate stereo and texture information for judgments of surface slant? Vision Res 43, 2539–2558 (2003).

55. Fleming, S. M. Metacognition and Confidence: A Review and Synthesis. Annu Rev Psychol 75, 241–268 (2024).

56. Zheng, Y., Recht, S. & Rahnev, D. Common computations for metacognition and meta- metacognition. Neurosci Conscious 2023, (2023).

57. Zheng, Y., Xue, K., Shekhar, M. & Rahnev, D. Similar computational noise for perceptual decision making with confidence, expectation, and reward. Preprint at 10.31234/osf.io/ydx6z (2024).

58. Yeon, J., Shekhar, M. & Rahnev, D. Overlapping and unique neural circuits are activated during perceptual decision making and confidence. Sci Rep 10, 20761 (2020).

59. Faivre, N., Filevich, E., Solovey, G., Kühn, S. & Blanke, O. Behavioral, Modeling, and Electrophysiological Evidence for Supramodality in Human Metacognition. The Journal of Neuroscience 38, 263–277 (2018).

60. Mazancieux, A., Fleming, S. M., Souchay, C. & Moulin, C. J. A. Is there a G factor for metacognition? Correlations in retrospective metacognitive sensitivity across tasks. J Exp Psychol Gen 149, 1788–1799 (2020).

61. Bruns, P. The Ventriloquist Illusion as a Tool to Study Multisensory Processing: An Update. Front Integr Neurosci 13, (2019).

62. Maiworm, M. & Röder, B. Suboptimal auditory dominance in audiovisual integration of temporalcues. Tsinghua Sci Technol 16, 121–132 (2011).

63. Plaisier, M. A., van Dam, L. C. J., Glowania, C. & Ernst, M. O. Exploration mode affects visuohaptic integration of surface orientation. J Vis 14, 22–22 (2014).

64. Rosas, P., Wagemans, J., Ernst, M. O. & Wichmann, F. A. Texture and haptic cues in slant discrimination: reliability-based cue weighting without statistically optimal cue combination. Journal of the Optical Society of America A 22, 801 (2005).

65. Bertana, A., Chetverikov, A., van Bergen, R. S., Ling, S. & Jehee, J. F. M. Dual strategies in human confidence judgments. J Vis 21, 21 (2021).

66. Macmillan, N. & Creelman, D. Detection Theory: A User’s Guide. (Psychology Press, 2004).

67. Noppeney, U. Perceptual Inference, Learning, and Attention in a Multisensory World. Annu Rev Neurosci 44, 449–473 (2021).

68. Rohe, T. & Noppeney, U. Cortical Hierarchies Perform Bayesian Causal Inference in Multisensory Perception. PLoS Biol 13, e1002073 (2015).

69. Rohe, T. & Noppeney, U. Distinct Computational Principles Govern Multisensory Integration in Primary Sensory and Association Cortices. Current Biology 26, 509–514 (2016).

70. Guipponi, O. et al. Multimodal Convergence within the Intraparietal Sulcus of the Macaque Monkey. The Journal of Neuroscience 33, 4128–4139 (2013).

71. Kayser, S. J., Philiastides, M. G. & Kayser, C. Sounds facilitate visual motion discrimination via the enhancement of late occipital visual representations. Neuroimage 148, 31–41 (2017).

72. Alink, A., Singer, W. & Muckli, L. Capture of Auditory Motion by Vision Is Represented by an Activation Shift from Auditory to Visual Motion Cortex. The Journal of Neuroscience 28, 2690–2697 (2008).

73. Herce Castañón, S., et al. Human noise blindness drives suboptimal cognitive inference. Nat Commun 10, 1719 (2019).

74. Maniscalco, B., Peters, M. A. K. & Lau, H. Heuristic use of perceptual evidence leads to dissociation between performance and metacognitive sensitivity. Atten Percept Psychophys 78, 923–937 (2016).

75. Boldt, A., Sun, Y. & Desender, K. Dis-confirmatory evidence drives confidence. Preprint at 10.31234/osf.io/tsr9z (2023).

76. Boundy-Singer, Z. M., Ziemba, C. M. & Goris, R. L. T. Confidence reflects a noisy decision reliability estimate. Nat Hum Behav 7, 142–154 (2022).

77. Winter, C. J. & Peters, M. A. K. Variance misperception under skewed empirical noise statistics explains overconfidence in the visual periphery. Atten Percept Psychophys 84, 161–178 (2022).

